# VOCAL SIMILARITY AND DIFFERENCES IN FOUR SPECIES OF THE GENUS *ANTHUS*: ACOUSTIC FEATURE ANALYSIS OF SOME COMMON CALLS

**DOI:** 10.1101/2022.09.25.509367

**Authors:** Marco Dragonetti

**Affiliations:** Gruppo Ornitologico Maremmano, c/o Museo di Storia Naturale di Grosseto, Strada Corsini 5, 58100, Grosseto, Italy

**Keywords:** call similarity, acoustic recognition, genus *Anthus*, vocal behaviour, taxonomy

## Abstract

Birds of the same genus often share similar call repertoires, the aim of this paper is to find species-specific and common acoustic features across species. This can be a useful tool for identification purposes and for studying intra-interspecific communication. Similar flight call (*tsip*) in two closely related species (*Anthus pratensis, Anthus spinoletta*) were studied to find characteristics, that allows to discriminate the two species with acoustic means. Three different call types (*tsip, soft* and *alarm*) of four species of the genus Anthus (*A. pratensis, A. spinoletta, A. petrosus, A. cervinus*) were also studied to find whether these common call types show different degrees of similarity. Discriminant function analysis correctly classified 98.4% of *A. pratensis* and *A. spinoletta* flight calls. Three acoustic parameters showed the highest discrimination power: the frequency modulations, the maximum frequency value and the minimum frequency value of the peak frequency contour. Using these three values I proposed a simpler procedure for recognizing these two species, that allowed a correct classification of 96% of calls.The three call types of the four *Anthus* species were studied using cross correlation among spectrogram contours. *Alarm* calls of the four species showed stronger similarity, while the other call types were more distinctive, with *soft* call seeming to have a lower similarity between species and hence a higher distinctive power. These results suggest the hypothesis that *alarm* call is similar, because it retains features of a common ancestor easing heterospecific communication, while the other calls showed decreasing similarity and more species-specific features.

## INTRODUCTION

Investigating the acoustic features of the vocalisations among species of the same genus may help to better understand their communication strategies and to shed light about the variable acoustic structures across related taxa. Moreover, acoustic analysis can be an useful tool for identification of morphologically similar species. Common call types used all year round offer a fruitful field of investigation for these purposes.

Meadow pipits (*Anthus pratensis*) and Water pipits (*Anthus spinoletta*) are morphologically similar species, they have partly overlapping wintering range and often share feeding areas (Cramp 1983; Alström and Mild 2003; Brichetti and Fracasso 2007). These congeneric similar species cope with problems of intraspecific and interspecific communication, therefore they should have finely tuned capability of call discrimination between species as found by Elfström (1992) for *A. pratensis* and *A. petrosus*, but in some cases they might also benefit from interspecific communication, e.g. in mixed flock with contact calls and with common danger signals. Outside the breeding season the vocal repertoire of these two species rely mainly on two call types, a *tsip* call used more often in flight and a *soft* call typically produced between members of a flock as a contact signal (Elfström 1992; Alström and Mild 2003). The former shows a very similar spectrographic structure for both species (Alström and Mild 2003; Bergmann et al. 2008), although it can sound slightly different for a trained ear (Brichetti and Fracasso 2007).

Elfström (1992) showed that calls of *A. pratensis* and *A. petrosus* contain species-specific characteristics and that these species are able to discriminate between conspecific and heterospecific calls of similar configuration at a distance. Finding comparable species-specific differences also for *tsip* call of *A. pratensis* and *A. spinoletta* would support the hypothesis that this call type is mainly used for intraspecific communication.

In addition, finding species-specific features of a common flight call can enable vocal identification of species that have similar winter plumage and behaviour. This can be a useful tool for field work when the two sister species share the same habitat and in the case of passive recording of migratory birds. Indeed, flight call recording and identification of nocturnal migrant birds is a method increasingly used (Hamilton 1962; Gal 2003; Farnsworth 2005). Some evidence suggest that birds of the genus *Anthus*, although mainly diurnal migrants (Elkins 2005), can migrate also by night (Ebenhöh 1998; Gal 2003; Wang 2019; Briedis et al. 2020), therefore acoustic identification turns to be interesting for these species too.

For years bird calls were considered innate (see for example Hamilton 1962), because subjects raised away from their own species developed abnormal songs but normal calls. After the discovery of developmental plasticity in the calls of several bird species (Mudinger 1970; Mudinger 1979; Marler and Slabbekoorn 2004), learning appeared to play a role in the call repertoire of some passerines. Particularly flight calls of cardueline finches showed remarkable plasticity and dialects were discovered for other call types (Baker et al. 2000; Marler and Slabbekoorn 2004). Although bird calls have not been studied as extensively as bird song, today it is commonly accepted that some calls are learned while others appear to be innate (Zann 1985; Williams 2008). Some calls can share very similar characteristics in several species of related taxa (Stefanski and Falls 1972; Fallow et al. 2011; Gayk et. al 2021), while others can be more or less distinctive for intraspecific recognition and/or communication (Elfström 1992; Kroodsma 2000; Fijen 2014). Genetic relatedness should favour a stronger similarity mainly for those innate calls that convey crucial information involved with immediate issues of life and death. On the other hand there are also calls, either genetically determined or developed by learning processes, which contains distinctive features and are important for species-specific recognition (Nowicki 1989, Marler and Slabbekoorn 2004).

In this paper I have investigated the acoustic features of a common flight call (*tsip*) of two sister species (*A. pratensis, A. spinoletta*). The aim is to find whether there are objective measurable differences among vocal parameters of *tsip* call, which is fairly similar for these two species. If consistent quantitative differences are found, it is interesting to find a measurable vocal threshold that allows a safe and relatively easy identification of the species. Finding quantitative differences for one call type poses the question whether this pattern is also found for other common call types in closely related species of the genus *Anthus*.

Interestingly, the acoustic similarity of some call types in different species can be the result of retaining features from a common ancestor (De Kort and Ten Cate 2001) or convergence in acoustic structure on the most effective design for a particular environment (Marler 1955; Marler and Slabbekoorn 2004). Broadening the view on other species of the genus *Anthus* and investigating the acoustic similarity and differences of different call types seemed me an useful approach to shed some light on these topics. Therefore three different call types (*tsip, soft* and *alarm*) of four species (*A. pratensis, A. spinoletta, A. petrosus, A. cervinus*) were studied. The results of this analysis, although have to be considered preliminar given the limitations of the dataset, can be useful for the study of intra/interspecific vocal communication and genetic relatedness among sister species of the genus *Anthus*.

## METHODS

I recorded calls of *Anthus pratensis* and *Anthus spinoletta* in central Italy with a Fostex FR2 digital recorder in wav uncompressed format (48 kHz sampling rate, 16 bit resolution). The microphone was a Sennheiser MKH20 in the focus of a 22” Telinga parabolic dish. The species identification was made visually (with binoculars, cameras and spotting scopes) before starting the recording session. Moreover almost all recordings of *A. spinoletta* were made in nesting areas of this species, where *A. pratensis* is absent.

I made 34 good quality recordings of *A. pratensis* and 31 of *A. spinoletta* from 2004 to 2021 in different places of Central Italy. The minimum distance between sampling places for each species, when recordings were made during the same season, was 25 km; the minimum time gap, when recordings were made in the same zone, was 26 months, therefore it is very unlikely that the same birds were recorded more than once, given that both species are not resident in Central Italy. The average of the acoustic parameters of all good quality calls was calculated from each recording, yelding a dataset of 65 independent measures. The *tsip* call type (in flight) was selected from these recordings following the classification and spectrographic representation available in the literature (Cramp 1983; Elfström 1992; Alström and Mild 2003; Bergmann 2008; Fijen 2014; Garner et al. 2015). All spectrograms were made with Raven Pro 1.6.1 (K. Lisa Yang Center for Conservation Bioacoustics 2019): window and DFT size 1024 samples, overlap 75%, window type Hann (3dB filter bandwidth 67.5 Hz).

One *tsip* call template of the two species (Fig. 1) was chosen among good quality recordings, with a signal to noise ratio above 10 calculated on the amplitude values expressed by Raven in arbitrary units. This call type has at least three visible harmonics and the second is always the dominant one, therefore a bandpass filter was used for limiting the analysis to this harmonic. As shown in Figure 1 most sound energy of the template *tsip* calls is concentrated between 4 and 8.5 kHz, for this reason I used the band limited energy detector of Raven Pro to select all the *tsip* calls to be analysed. The detector parameters were: minimum frequency 4000 Hz, maximum frequency 8500 Hz, minimum duration 0.03 s, maximum duration 0.2 s, minimum separation 0.02 s, minimum occupancy 50%, SNR (signal to noise ratio) threshold 5 dB. The detections showing unwanted signals (e.g. overlapped calls of the target species or of other species, various noises) were discarded. A detailed acoustic analysis was performed on 212 *A. pratensis* and 165 *A. spinoletta tsip* calls and data were averaged for each recording, yielding 34 and 31 independent data for the two species; the following parameters were calculated by means of the appropriate Raven Pro functions: Center Frequency (CF), Frequency Contour Percentile 75 (FC75), Peak Frequency Contour Max Frequency (PFCMa), Peak Frequency Contour Min Frequency (PFCMi), Duration 90% (DUR90). The difference between PFCMa and PFCMi was calculated as a measure of the frequency bandwidth (BW). I counted for each call the number of relative maximum plus minimum values (RMM) found by the FC75 function. The time interval between the beginnings of consecutive call selections (TI) was measured. In 2020/2021 further 5 recordings of *A. pratensis tsip* calls were made in Northern and Central Italy and other 5 recordings of *A. spinoletta tsip* calls from Central Europe and Northern Italy were provided by Xeno-Canto recordists in wav uncompressed original format. These 10 recordings (for a total of 45 calls) were used to test the ability of DFA to discriminate the two species.

**Figure 1.**
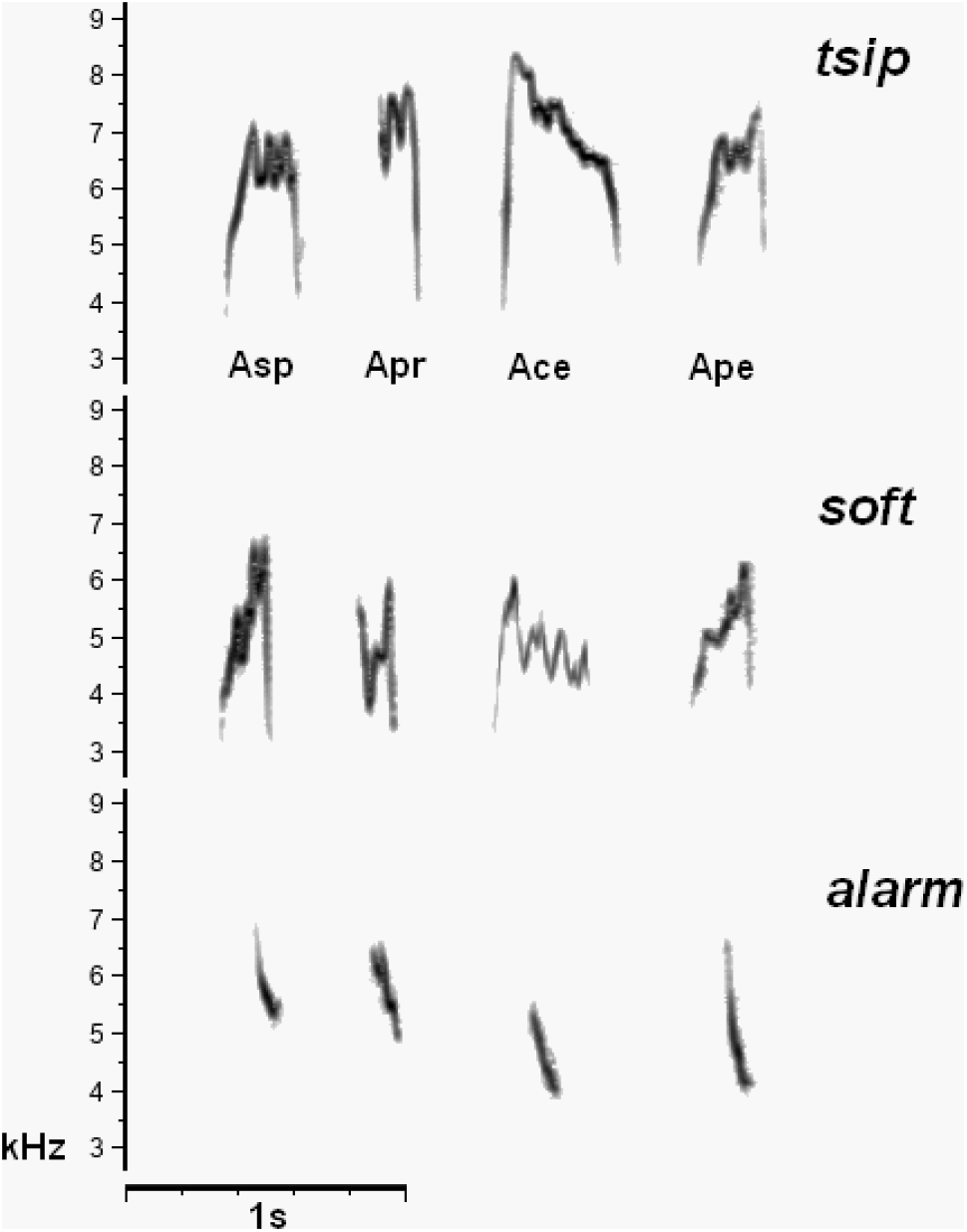
Spectrograms of three different call types of *A. spinoletta* (Asp), *A. pratensis* (Apr), *A. cervinus* (Ace), *A. petrosus* (Ape). See Methods for call type classification

The three different call types (*tsip, soft* and *alarm*) of the four *Anthus* species (*A. pratensis, A. spinoletta, A. petrosus, A. cervinus*) were compared following the classification and spectrographic representation of Elfström (1992) and Alström and Mild (2003). Figure 1 shows the template calls chosen for each call type of the four species. 13 recordings of *tsip* call and 12 recordings of *soft* calls for *A. pratensis* and *A. spinoletta* were randomly selected from the dataset above described, while 13 recordings of *tsip* call for *A. cervinus* and *A. petrosus*, 12 recordings of *soft* call for *A. cervinus* and *A. petrosus* and 10 recordings of *alarm* call for *A. cervinus, A. petrosus, A. pratensis* were audio files published on Xeno-Canto (randomly selected among all those available), 10 recordings of *alarm* call for *A. spinoletta* were made in Central Italy by the author as specified above. Unfortunately most of Xeno-Canto recordings were not available in an original non-compressed (wav) format, but only as mp3 files. Since all the recordings were not homogeneous as file format, bitrate, recording tools and editing processes, a detailed analysis of all the acoustic parameters, as made for *A. pratensis* and *A. spinoletta tsip* call, could have yielded unreliable results. To avoid this drawback I preliminarily tested the capability of Raven Pro FC75 function to assess in a sufficiently precise manner general spectral and temporal characteristics of the same calls recorded in three different ways: wav original uncompressed format, mp3 compressed file of medium quality (170/210 kbps), mp3 compressed format of low quality (145/185 kbps). The results of this test (shown in the supplementary material) suggested that FC75 function was able to assess correctly general bioacoustic characteristics for all the different quality levels. Similarly Araya-Salas et al.(2019) found that compression did not significantly bias the acoustic measurements of similarity. Therefore the comparison among the three call types in the four *Anthus* species was made using only the FC75 parameter. From each recording of *alarm* call 7 to 9 call contours were averaged yelding 10 independent data for each *Anthus* species, from each recording of *tsip* call 4 to 8 call contours were averaged yelding 13 data, from each recording of *soft* call 3 to 9 call contours were averaged yelding 12 data. The total number of contours averaged for each call type and for each species was always 84. The call selection was made with the same method described above, using the band limited energy detector of Raven Pro (band limits: 4000 - 8500 Hz, 3000 - 7000 Hz, 3500 - 7000 Hz for *tsip, soft* and *alarm* call respectively). All recordings were filtered with a bandpass filter for limiting the analysis to the dominant harmonic where most of sound energy was found.

### Statistical analysis

Where not specified otherwise all the statistical tests were performed using PAST v. 4.09 package (Hammer et al. 2001). Multivariate normality of data was checked with omnibus test by Doornik and Hansen (2008). The comparison of *A. pratensis* versus *A. spinoletta tsip* call was performed with PERMANOVA based on Euclidean distance measure (Anderson 2001). Principal components analysis (PCA) was used to find hypothetical variables (components) accounting for as much as possible of the variance in these multivariate data (Legendre and Legendre 1998). The PCA and PERMANOVA were performed including all the seven varaiables shown in Table 1. Coefficient of variation (CV) was calculated with the following formula: CV =100*(1+1/4n)*sd/m (Scherrer 1984; Aubin et al. 2004), where n = number of observations, SD = standard deviation, m = mean. I have also calculated a CVr as the ratio of interspecific (CVis) versus average intraspecific coefficient of variation (CVin). CVr is a measure of the identification power of each vocal parameter, if it is higher than 1 the interspecific differences are heavier than the intraspecific ones and of course the higher is CVr value the better is the discrimination power of the parameter. The equivalence of the variance/covariance on multivariate samples was checked with Box’s M test (Rencher 2002); this is a prerequisite to perform the discriminant function analysis (DFA), used to find the differences among acoustic parameters of *A. pratensis* versus *A. spinoletta tsip* calls.

**Table 1.**
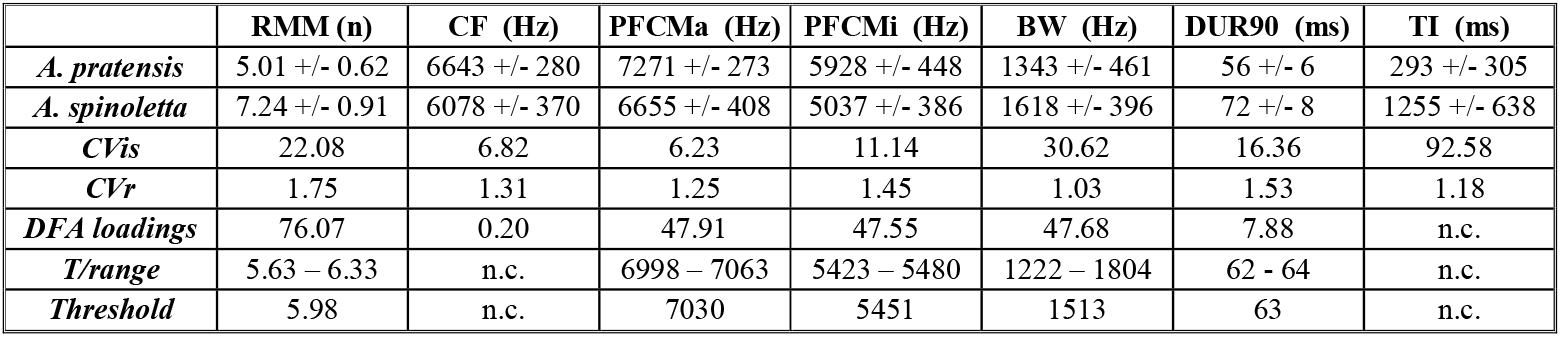
Comparison of *tsip* call vocal parameters between *A. pratensis* and *A. spinoletta*. **CV** = coefficient of variation (see Methods); **CVis** = interspecific coefficient of variation; **CVr** = ratio between CVis and CV intraspecific of the two species. **DFA** = discriminant function analysis. **T/range** = range between mean + SD of lower value parameter and mean - SD of higher value parameter. **Threshold** = median of T/range values taken as separation threshold between the two species. **RMM** = number of relative minimum and maximum values of call frequency contour; **CF** = center frequency; **PFCMa** = maximum of the peak frequency contour measured from spectrogram slices; **PFCMi** = minimum of the peak frequency contour; **BW** = call frequency bandwidth calculated from (PFCMa - PFCMi); **DUR90** = call duration containing 90% of the call energy; **TI** = time interval between start of subsequent calls; n.c. = not calculated. *A. pratensis* n=34, *A. spinoletta* n=31

|  | **RMM (n)** | **CF (Hz)** | **PFCMa (Hz)** | **PFCMi (Hz)** | **BW (Hz)** | **DUR90 (ms)** | **TI (ms)** |
| --- | --- | --- | --- | --- | --- | --- | --- |
| *A. pratensis* | 5.01 +/- 0.62 | 6643 +/- 280 | 7271 +/- 273 | 5928 +/- 448 | 1343 +/- 461 | 56 +/- 6 | 293 +/- 305 |
| *A. spinoletta* | 7.24 +/- 0.91 | 6078 +/- 370 | 6655 +/- 408 | 5037 +/- 386 | 1618 +/- 396 | 72 +/- 8 | 1255 +/- 638 |
| **CV<sub>is</sub>** | 22.08 | 6.82 | 6.23 | 11.14 | 30.62 | 16.36 | 92.58 |
| **CV<sub>r</sub>** | 1.75 | 1.31 | 1.25 | 1.45 | 1.03 | 1.53 | 1.18 |
| **DFA loadings** | 76.07 | 0.20 | 47.91 | 47.55 | 47.68 | 7.88 | n.c. |
| **T/range** | 5.63 – 6.33 | n.c. | 6998 – 7063 | 5423 – 5480 | 1222 – 1804 | 62 - 64 | n.c. |
| **Threshold** | 5.98 | n.c. | 7030 | 5451 | 1513 | 63 | n.c. |

The comparison among four *Anthus* species (*A. pratensis, A. cervinus, A. spinoletta, A. petrosus*) was performed using FC75 to find similarity and differences in the three different call types (*alarm, tsip, soft*). The randomly selected call contours were averaged for each recording and an averaged contour was then calculated for the four species in the three different call types. Each averaged contour represented a series of frequency data evenly spaced in time (5 ms apart). Mantel correlogram (Legendre and Legendre 1998), that is a multivariate extension to autocorrelation based on any similarity or distance measure, made a first comparison among all four species contours of each call type. The correlation index of Mantel test showed the similarity between the averaged call contours shown in Figure 3. A second approach was made with a more classical cross-correlation comparing the time series obtained with the FC75 function. All paired comparisons were performed between average call contours from every recording of the four species for each different call type with the following constraints: each recording must be used only once, all recordings must be used, each pair of recording comparison was randomly chosen. This scheme yielded for example 10 comparisons for *alarm* call of *A. pratensis* versus *A. spinoletta*, 10 comparisons of *A. pratensis* versus *A. cervinus* …and so on. These values of cross correlation were averaged for each pair of comparisons among the four species and for each call type. This way it was obtained a similarity value for each possible comparison among the four species and a mean similarity value for all the comparisons among the four species for each call type (Table 1 row 8). The average correlation index was calculated at two different lags (lag 0 and lag m of maximum correlation index); the average value of the lag accounting the maximum correlation index was also calculated. Of course a higher correlation index at lag 0 or near 0 expresses a higher similarity among different species for that particular call type. The cross-correlation was calculated using the R function ccf (R core Team 2020).

## RESULTS

### Comparison between *tsip* call of *A. pratensis* versus *A. spinoletta*

Table 1 shows *tsip* call parameters of *A. pratensis* (n= 34; each data is average of 3 to 10 calls, mean 6.2) and *A. spinoletta* (n= 31; each data is average of 2 to 9 calls, mean 5.3). Doornik and Hansen (2008) omnibus test indicates a highly significant (p<0.001) departure from normality of these *tsip* call data. Therefore the two species were compared by means of non parametric PERMANOVA test, which showed a highly significant difference (F=45; P<0.001) of acoustic features.

Principal component analysis (PCA) of all multivariate data (Figure 2) was performed. The three first components accounted for 89.4% of the variance and PCA loadings of the variables showed values ranging between 0.33 (DUR90 and RMM), 0.34 (BW) and 0.38 (CF, PCFMa, PCFMi), but TI showed a lower value (0.28). Since TI has several missing data and also a very high CVin (= 78.5) with a quite low CVr (Table 1), it was excluded from further analyses because of its unreliability.

**Figure 2.**
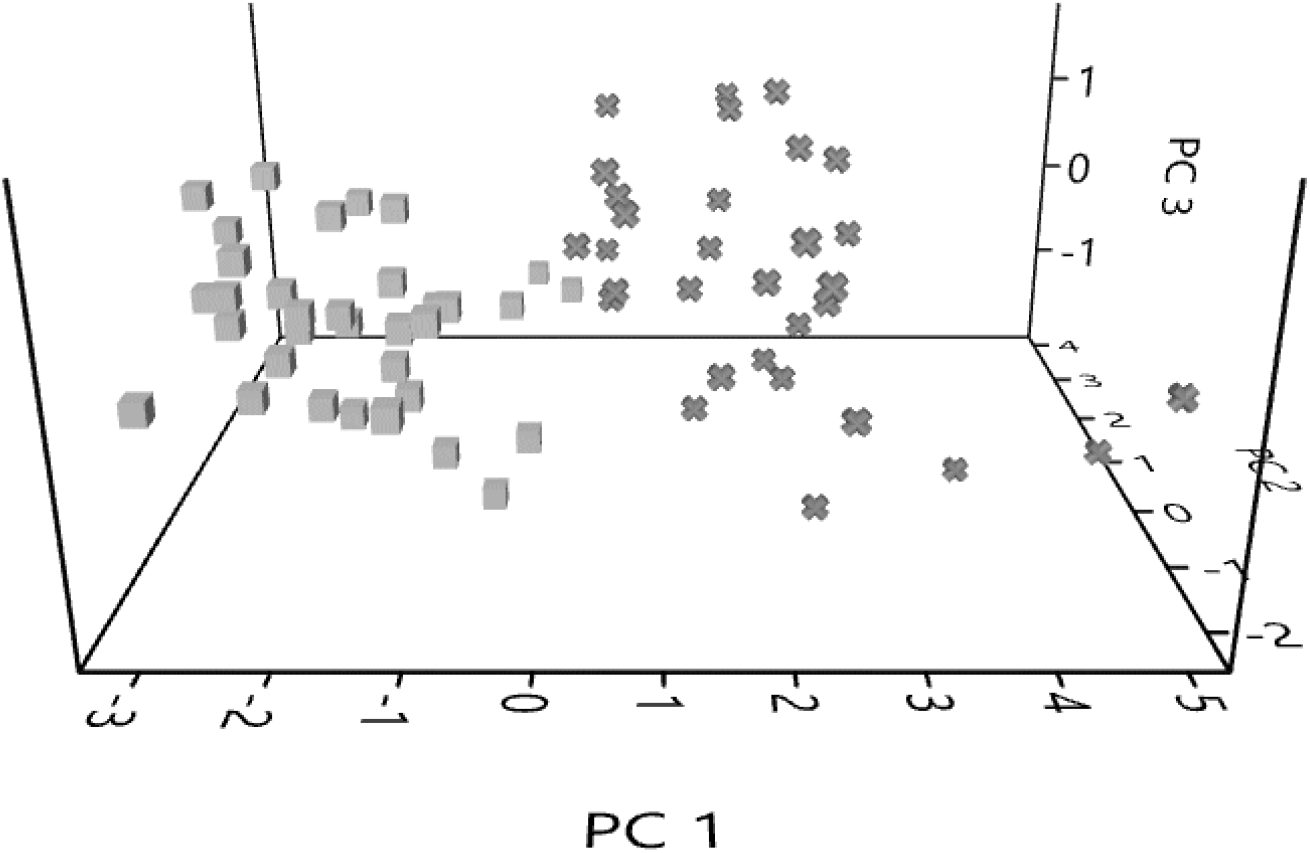
Comparison of *tsip* call vocal parameters between *A. pratensis* (cubes) and *A. spinoletta* (crosses). Scatter plot in the coordinate system given by the first three PCA components that account for the 89.4 % of the variance.

**Figure 3.**
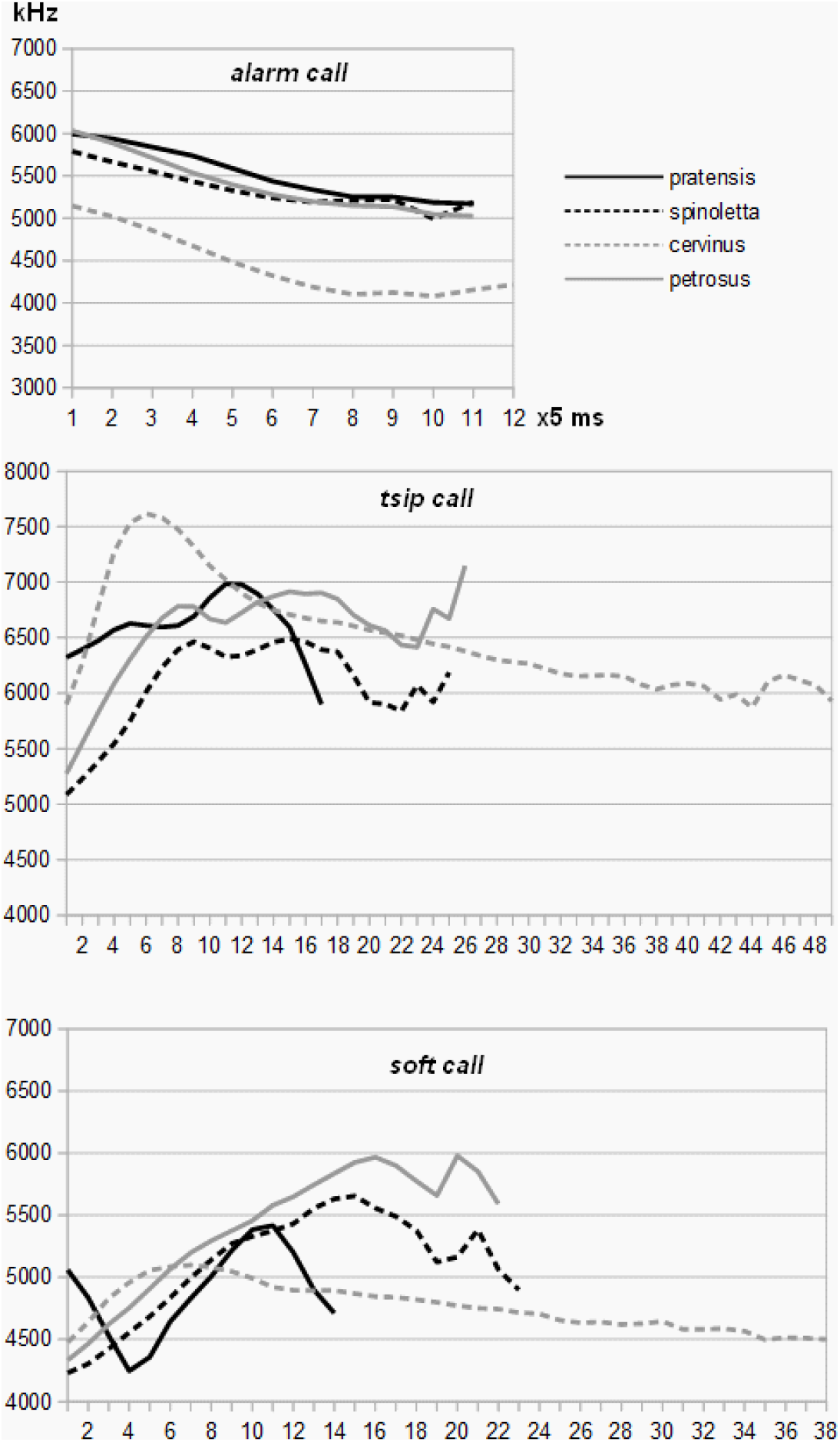
Average frequency contours (FCP75, see Methods) of three different calls in four *Anthus* species. ***Alarm* call**: average of 10 recordings for each species. ***Tsip* call**: average of 13 recordings for each species. ***Soft* call** : average of 12 recordings for each species. The total number of randomly selected calls for every call type of each species is always 84. Time series (x axis) are made by points evenly spaced 5 ms apart

CVr was definitely higher than 1 for all acoustic parameters (except BW that had the lower value of 1.03, Table 1) showing a good potential predictor power of these acoustic features for the recognition of the two sister species.

The Box’s M test (Rencher 2002) for equality of variance-covariance of multivariate data (excluding TI parameter) yielded a not significant result (F=1,38). Moreover the normality test (Doornik and Hansen 2008) showed a slightly significant value (p=0.045), accounting for a slight departure from normality of the data. Hence the prerequisites to perform DFA were satisfied, since Sever et al. (2005) showed the DFA to be highly robust to non-normal data.

DFA correctly classified 98.4% of calls with jack-knifed validation. Only one *A. pratensis* recording was misclassified as *A. spinoletta*. Further 5 recordings of *A. spinoletta* and 5 of *A. pratensis*, included in the same discriminant function as unknown specimens, were all correctly classified. The acoustic parameters with higher DFA loadings, accounting for a better discriminant power, were RMM, PCFMa, PCFMi and BW (Table 1). DUR90 has a very low loading and CF a near zero value.

I have calculated threshold values that can discriminate the *tsip* call of the two species (Table 1), as median of range between mean + SD of lower value parameter and mean - SD of higher value parameter. I tested the acoustic data using threshold values of three parameters (RMM, PCFMa and PCFMi) with the following rule: i) at least two parameters must be in agreement; ii) one of the two must be the RMM. This procedure was able to classify correctly 96% of all available recordings (n=75).

### Similarity and differences among four *Anthus* species calls

Figure 3 shows the average contours of *alarm, tsip* and *soft* calls for *A. spinoletta, A. pratensis, A. cervinus* and *A. petrosus*. An objective measure of similarities among species for each different call type was calculated with two approaches. A multivariate Mantel correlogram, computed with correlation index, was applied on the averaged time series of Figure 3. This test showed a high similarity of *alarm* call of the four species (0.981) and lower values for *tsip* (0.660) and *soft* calls (0.421).

Table 2 shows the results of a classical cross-correlation applied to the six possible comparisons between the four species for each of the three different call type. Average value of *alarm* call is again by far higher (0.943) than *tsip* and *soft* calls (0.441 and 0.329). Cross-correlation indexes, either at lag(0) or lag(m), were not normally distributed and variance-covariance was significantly unequal. Therefore I performed the comparisons among the three call types with Kruskall-Wallis test. Cross-correlation indexes, both at lag (0) and at lag (m) of maximum correlation (Table 2), showed a significant difference among the three call types (Hc=75.9 p<0.001 at lag 0 and Hc=78.3 p<0.001 at lag m). Mann Whitney post hoc test (with continuity correction and a correction for ties) was significant for the comparisons *alarm* vs *tsip* call (p<0.001) and *alarm* vs *soft* call (p<0.001), while the comparison *tsip* vs *soft* call was not significant. The average lag of maximum correlation was 0.05 for *alarm* call, but shifted to 1.71 for *tsip* and to 3.31 for *soft* call. Again Kruskall-Wallis test showed a significant difference among these lag data (Hc=54.5 p<0.001). Mann Whitney post hoc test showed a significant shift of lag for *tsip* and *soft* versus *alarm* call (p<0.001), but also a significantly higher shift for *soft* versus *tsip* call (p<0.01). An outlier data was produced by the comparisons between *A. spinoletta* versus *A. petrosus* both for *tsip* and *soft* calls (Table 2), which yielded a maximum correlation coefficient of 0.889 and 0.860 at lag 0. These two data, although significantly lower than *alarm* call coefficient (p<0.05), were clearly by far higher than the other coefficients calculated for these two call types. Moreover, for *A. spinoletta* versus *A. petrosus* comparison, the average lag of maximum correlation (avg. lag m) did not shift significantly from zero also for *tsip* and *soft* calls.

**Table 2.** Cross correlation results (average +/-SD) among contours of three different call type (*alarm* call, *tsip* call, *soft* call) in four *Anthus* species; **pr =** *A. atensis*; **sp =** *A. spinoletta*; **ce =** *A. cervinus*; **pe =** *A. petrosus*. **Mean =** average of all the cross-correlations of all comparisons. **(0) =** cross-correlation results at lag **(m) =** cross-correlation results at lag of maximum correlation coefficient; **avg. lag(m) =** average lag value corresponding to the maximum correlation coefficient; **a) =** number of cross-correlations performed for *alarm* calls; **n(t) =** number of cross-correlations for *tsip* calls; **n(s) =** number of cross-correlations for *soft* calls. ch contour submitted to cross-correlation is an average of minimum 3 and maximum 9 randomly selected calls from each recording of a given species. The total mber of randomly selected calls is always 84 for each recording set and for every call type of the four species.

|  | n (a) | alarm(0) | alarm(m) | avg. lag (m) | n (t) | tsip(0) | tsip(m) | avg. lag (m) | n (s) | soft(0) | soft(m) | avg. lag (m) |
| --- | --- | --- | --- | --- | --- | --- | --- | --- | --- | --- | --- | --- |
| **pr vs sp** | 10 | 0.869+/-0.151 | 0.905+/-0.085 | 0.29+/-0.49 | 13 | 0.433+/-0.173 | 0.502+/-0.128 | 1.14+/-0.90 | 12 | 0.463+/-0.132 | 0.650+/-0.089 | 1.86+/-0.38 |
| **pr vs ce** | 10 | 0.962+/-0.021 | 0.962+/-0.021 | 0.00 | 13 | 0.313+/-0.196 | 0.447+/-0.070 | 2.71+/-3.15 | 12 | -0.06+/-0.394 | 0.711+/-0.140 | 3.71+/-1.38 |
| **pr vs pe** | 10 | 0.935+/-0.041 | 0.935+/-0.041 | 0.00 | 13 | 0.367+/-0.234 | 0.455+/-0.145 | 1.29+/-1.25 | 12 | 0.388+/-0.118 | 0.582+/-0.106 | 2.00+/-0.82 |
| **sp vs ce** | 10 | 0.963+/-0.045 | 0.963+/-0.045 | 0.00 | 13 | 0.318+/-0.194 | 0.462+/-0.085 | 2.43+/-1.40 | 12 | 0.175+/-0.450 | 0.481+/-0.182 | 4.86+/-3.76 |
| **sp vs pe** | 10 | 0.960+/-0.034 | 0.960+/-0.034 | 0.00 | 13 | 0.889+/-0.034 | 0.889+/-0.034 | 0.00 | 12 | 0.860+/-0.077 | 0.871+/-0.068 | 0.43+/-0.053 |
| **ce vs pe** | 10 | 0.968+/-0.027 | 0.968+/-0.027 | 0.00 | 13 | 0.325+/-0.131 | 0.383+/-0.061 | 2.71+/-4.03 | 12 | 0.146+/-0.371 | 0.437+/-0.212 | 7.00+/-3.37 |
| **Mean** | 60 | 0.943+/-0.074 | 0.949+/-0.049 | 0.05+/-0.22 | 78 | 0.441+/-0.263 | 0.523+/-0.192 | 1.71+/-2.34 | 72 | 0.329+/-0.407 | 0.622+/-0.198 | 3.31+/-3.00 |

## DISCUSSION

This paper showed that a similar flight call (*tsip*) of two sister species, *A. pratensis* and *A. spinoletta*, contains species-specific characteristics, that allows to discriminate the two species only with acoustic means. Some results of my research are similar to those reported by Elfström (1992) for *A*.*pratensis* and *A. petrosus*: for both studies, although the parameters are measured in different ways, *A. pratensis* has a lower call duration, a reduced frequency range (BW of Table1) and a less frequency modulated call, represented by RMM parameter in this paper and by pulse number in Elfström’s work (1992). The top frequency figure of *A*.*pratensis* (PFCMa Table 1) is quite consistent with that reported by Elfström (1992). Higher and lower frequencies of *A. spinoletta* are close to those reported by Fijen (2014). Multivariate analysis of all acoustic features showed a highly significant difference (Table 1) between the two species. The first question is: is it possible to reduce the parameters that are effective and reliable to discriminate the two species? The low loading value in the PCA analysis, the relatively low CVr (Table 1), the very high intraspecific variability (CVin = 78.5, that means a fairly unreliable parameter) and several missing data, lead to discard time interval between calls (TI). Frequency bandwidth (BW) proved to be less reliable because of its very low CVr (1.05) and high variability (CVin = 29.7). Therefore the acoustic parameters were now reduced to five (RMM, CF, PCFMa, PCFMi, DUR90).

The second question is: what is a suitable method to recognize acoustically the species and to classify new unknown data? DFA is a commonly used method in bioacoustics to classify multivariate parameters extracted from bird vocalizations (Galeotti and Sacchi 2001; Lengagne 2001; Christie et al. 2004; Terry *et al*. 2005) and it allows classifying new unknown specimens that are not part of the dataset used to create the discriminant function, so that future observations can be correctly grouped. Moreover DFA helps to understand what are the most important acoustic features to distinguish the two sister species. In this paper I found that the discriminant function is able to correctly classify 98.4% of the recordings and group assignment is validated by a leave-one-out cross-validation (jack-knifing) procedure. Only one bird out of 65 was misclassified. The DFA correctly classified all further 10 recordings of the two *Anthus* species that were added to the discriminant function as unknown specimens. An analysis of the DFA loadings (Table 1) shows that two parameters (central frequency CF and call duration DUR90) have very low values compared with the other ones, therefore their discrimination power is relatively low. Since BW has a good DFA loading, but, as stated above, it is less reliable than the other parameters, the most important acoustic features in the discrimination process are the remaining three parameters: RMM representing the frequency modulations of the tsip call, PFCMa that is the maximum frequency value of the peak frequency contour and PFCMi that is the minimum frequency value of the peak frequency contour. Among these last three parameters RMM shows the highest loading value for the DFA and can thus be considered the most critical acoustic feature for the recognition of these two bird species. I also tested a simpler and lighter procedure to discriminate the *tsip* calls of *A. pratensis* and *A. spinoletta*, which needs only the three above mentioned parameters. Table 1 at the sixth row shows for each acoustic parameter the lower and higher values yielded by the mean minus the SD and the mean plus SD: these ranges can reasonably used to find a threshold that separates the two species. The threshold for the three most important parameters (RMM, PFCMa and PFCMi) was calculated as the median of these ranges (Table 1). As shown above in the Results, these threshold values allowed a correct classification of 96% of recordings, where only 2 *A. pratensis* and 1 *A. spinoletta* were misclassified over 75 recordings. Of course either this procedure -that is relatively straitghtfoward and simpler than DFA - or the DFA itself may be successfully used only if the methods of acoustic analysis exposed in this paper are carefully followed.

This paper showed that two closely related species, with an overall similar vocalization, nevertheless possess fine, but significant differences of acoustic features that may be important to transmit species-specific information and to discriminate between conspecific and heterospecific vocalizations, like found by Elfström (1992) for *A. petrosus* and *A. pratensis*. This finding is confirmed by other authors for some species of the genus *Anthus* (Fijen 2014; Garner et al. 2015) and for bird species belonging to other taxa (Stiffler et al. 2018; Tietze et al. 2008). On the other hand, many bioacoustics researches found interspecific communication mediated by various bird call types (Johnson et al. 2003; Goodale and Kotagama 2005; Magrath et al. 2007; Templeton and Greene 2007; Magrath et al. 2009; Fallow et al. 2011; Fallow et al. 2013). The advantages of interspecific eavesdropping and communication by means of calls are documented in the literature (Nuechterlein 1981; Burger 1984; Forsmann and Mönkkönen 2001; Goodale and Kotagama 2005; Goodale et al. 2010) and this raises the question of how birds recognize heterospecific calls and their meaning. Learning has been suggested to explain avian recognition and response to heterospecific calls (Curio 1971; Hurd 1996; Magrath 2009), a second suggested hypothesis is that similar acoustic features facilitate heterospecific recognition (Marler 1957; Stefanski and Falls 1972; Johnson *et al*. 2003; Fallow *et al*. 2011) and a third hypothesis combines both previous ones: Hurd (1996) suggested that interspecific recognition results from both associative learning and common acoustic properties. Therefore it was useful to broaden my acoustic study to three different call types of four *Anthus* species, to study the heterospecific communication by means of some common call types in this genus.

The choice to analyse only the call contour (see Methods) was made for the following reasons: my aim was to represent objectively the overall acoustic features of the calls, underlining the trend of dominant frequency, which seems particularly involved in heterospecific recognition (Johnson et al. 2003); since most recordings were from Xeno-Canto mp3 files, with unknown quality level, I previously tested the reliability of contour representation (see Methods), that proved to be enough for the aims of the present work; finally, call representation by mean of average spectrogram contour is particularly suitable to be combined with cross-correlation as an objective method of evaluating vocal similarity. Combining cross-correlation with frequency contour is considered a method that may capture more precisely how signals vary over the duration of an acoustic unit (Odom *et al*. 2021).

The results of the comparison among *A. spinoletta, A. pratensis, A. cervinus* and *A. petrosus* on three different call types (*tsip, soft* and *alarm*) show that similarity of *alarm* calls in the four species is significantly and by far higher than the other two call types, with a tendency of a decrease in similarity from *tsip* to *soft* call. Indeed average correlation coefficient of the four species is significantly higher for *alarm* call either at lag 0 or at lag of the maximum correlation (Table 2). Since a shift from lag 0 proves a decreased similarity on the overall time shape of the calls, it is worth to analyse the average lag values where the correlation coefficient reach its maximum. For *alarm* it is 0.05, for *tsip* is 1.71 and for *soft* call is 3.31, where the differences of these values are highly significant, showing a decrease of similarity for the three call types among the four species. While the similarity among *alarm* calls of the four species is very strong and by far higher than the other two call types, the mean values of the correlation coefficients of *tsip* and *soft* calls don’t differ significantly, although the lag 0 values show a higher similarity for *tsip* than for *soft* call. This slight difference is confirmed by Mantel correlogram index: 0.660 for *tsip* and 0.421 for *soft* call. I can conclude that *alarm* calls of the four species show a very strong similarity, while the other two call types are more distinctive showing a much lower similarity, with *soft* call seeming to have a slightly lower similarity between species and hence a higher distinctive power. Since all four species are closely related from a taxonomic point of view, the results of my investigation support the hypothesis that *alarm* calls retain features of a common ancestor, while the other two call types (*tsip* and *soft*) show a decreasing similarity between the four *Anthus* species, resulting in a higher content of species-specific characteristics. The *alarm* call type studied in the present paper is only given in the vicinity of nest or in the breeding territory (Alström and Mild 2003; Bergmann et al. 2021; Dragonetti pers. obs.) therefore it is a very strong signal of danger, that the evolutive process may have maintained almost unaltered for these *Anthus* species, because it eases heterospecific communication. On the other hand *tsip* call is the most common call uttered in many different contexts mainly in flight and *soft* call is typically uttered between members of a flock as a contact signal (Elfström 1992; Alström and Mild 2003), therefore for these two calls the needs of intraspecific communication increase and evolution may have increased their acoustic divergence, as shown in the present paper.

Interestingly the comparison *A. petrosus* versus *A. spinoletta* yields a somewhat outlier data (see Table 2 row 5): indeed both *tsip* and *soft* calls show values (0.889 and 0.860 respectively) very close to that of *alarm* call (0.960) and a nearly null shift from lag 0, accounting for a greater similarity of this couple of species than the other ones. The taxonomy of “Water Pipit complex” has been much debated and until recently *A. petrosu*s and *A. spinoletta* were treated as a single species (Ali and Ripley 1998). Arctander et al. (1996) found only 1.2 % difference in mitochondrial cytochrome b sequences between *A. spinoletta* and *A. petrosus*, for this reason they expressed their reluctance to treat them as separate species. Today they are commonly treated as separate species, but there is no doubt that they are more closely related from a genetic point of view than the other species studied in this paper (see Voelker 1999) and this may explain the higher similarity indexes reported in Table 2.

A further consideration can be made about *alarm* call similarity in the genus *Anthus*, checking the eighteen species living in Europe, Asia and North America and visually comparing specimen spectrograms of *alarm* call available in the literature (Cramp and Simmons 1983; Alström and Mild 2003; Pieplow 2019; Bergmann et al. 2021). The *alarm* calls of six species were unknown (*A. nilghiriensis, A. silvanus, A. spragueii, A. rufulus, A. similis, A. campestris*) the remaining twelve might be classified as follows: three (*A. berthelotii, A. godlewskii, A. richardi*) show a completely different *alarm* call compared to those studied in this paper, seven (*A. trivialis, A. gustavi, A. roseatus, A. cervinus, A. petrosus, A. spinoletta, A. pratensis*) show a highly similar call and two (*A. hodgsoni, A. rubescens*) show a lower degree of similarity but are very reminiscent of the *alarm* described in this paper. If we look at the maximum likelihood tree of the genus *Anthus* based on the mitochondrial cytochrome b gene (from Voelker 1999), we can find all the former three species in the 4th group (the “large” pipits), while the remaining nine are all in the 3rd group (the “small” pipits). The results of the present paper support the hypothesis that *alarm* call of four *Anthus* species is highly similar, because it retains features of a common ancestor and may ease heterospecific communication. The call similarities of closely related species can be important for phylogenetic comparative studies, as suggested by the comparisons made in the *Anthus* genus. The method commonly used in phylogenetic studies is integrative taxonomy (combination and integration of multiple types of evidence, see Sangster 2018): together with genetic and morphological information, acoustic data play a major role in taxonomic designation of birds. The song of passerines, considered to be one of the most important traits promoting differentiation among species (Price 2008), was extensively investigated in phylogenetic comparative studies, but the data I reported here suggest that also the comparisons of call repertoires may be a useful tool for taxonomic studies.

## Supporting information

Supplemental results

## ACKNOWLEDGEMENTS

I am extremely grateful to Martin Boesman, Fernand Deroussen, Ivan Farronato, Niels Krabbe, Lars Lachmann, Dimitri Marrone, Nicolas Martinez, Jarek Matusiak, Marco Pesente, Maurizio Sighele, Bodo Sonnenburg, Stanislav Wroza who provided original recordings (either in uncompressed or compressed format) of *Anthus* species.Without their help this work could not be done. The Xeno-Canto collection was an invaluable resource for my work. The credits for all recordings I have used from Xeno-Canto platform can be accessed in the supplementary material. I am indebted to Gianni Pavan and Orlando Tommasini for their comments and suggestions on a first draft of the manuscript. I also thank two anonymous reviewers for their helpful comments on the manuscript.

