## Supplemental table 1 for "VOCAL SIMILARITY AND DIFFERENCES IN FOUR SPECIES OF THE GENUS *ANTHUS*: ACOUSTIC FEATURE ANALYSIS OF SOME COMMON CALLS"

***Tsip call A. cervinus***

| XC Recording | Recordist | Country | Location |
|---|---|---|---|
| 432617 | Lars Edenius | Sweden | Degernas |
| 435092 | Michèle Peron | Germany | Oberbayern |
| 462622 | Boris Delahaie | Morocco | Bir Anzerane road |
| 502377 | Jarek Matusiak | Poland | Lubelskie |
| 592483 | Alain Hofmans | Netherlands | Noord-Brabant |
| 597257 | Testaert Dominique | France | Nouv.Aquitaine |
| 603155 | Ewa Szczepankiewicz | Poland | Mazowieckie |
| 641550 | Lonnie Bregman | Netherlands | Frieseland |
| 644821 | Guillaume Petitjean | France | Bourgogne |
| 654154 | Oscar Campbell | Un. Arab Emirates | Fujairah |
| 670881 | Lars Edenius | Sweden | Umea |
| 673089 | Hannu Varkki | Finland | Uusimaa |
| 676385 | Toy Janssen | Netherlands | Noord-Brabant |

***Tsip call A. petrosus***

| XC Recording | Recordist | Country | Location |
|---|---|---|---|
| 112460 | Sander Bot | Netherlands | Frieseland |
| 146029 | Julien Rochefort | France | Brittany |
| 281353 | Peter Boesman | Belgium | W.-Vlaanderen |
| 284623 | Jarek Matusiak | Poland | Pomeranian Voiv. |
| 305656 | David Darrel-Lambert | United Kingdom | Kent |
| 345757 | Jarek Matusiak | Poland | Puck County |
| 368723 | Thimo Schnabel | France | Bretagne |
| 437785 | Jarek Matusiak | Poland | Pomeranian Voiv. |
| 443926 | David Pennington | United Kingdom | S. Yorkshire |
| 449678 | Patrik Aberg | Sweden | Gotland |
| 456942 | David Darrel-Lambert | United Kingdom | Essex |
| 522465 | Jarek Matusiak | Poland | Zatoka Gdanska |
| 687454 | Irish Wildlife Sounds | Ireland | County Donegal |

***Soft call A. cervinus***

| XC Recording | Recordist | Country | Location |
|---|---|---|---|
| 67213 | Patrik Aberg | Sweden | Vastergotland |
| 388003 | Niels Van Doninck | Belgium | Vlaanderen |
| 435076 | Michael Heiss | Azerbaijan | Besh Barmag |
| 147147 | Albert Lastukhin | Russian Federation | Chuvashia |
| 322598 | Thomas Luthi | Israel | Be'er Sheva |
| 342802 | Lars Buckx | Netherlands | Parnassia |
| 386519 | Sonnenburg | Germany | Brandenburg |
| 596357 | Bram Piot | Laos | Vientiane |
| 446999 | David Darrel-Lambert | Saudi Arabia | Al Madinah |
| 471999 | Stanislas Wroza | France | Haute-Corse |
| 537940 | Mathieu André | Belgium | Wallonie |
| 586250 | Lars Edenius | Sweden | Umea |

***Soft* call *A. petrosus***

| XC Recording | Recordist | Country | Location |
|---|---|---|---|
| 281352 | Peter Boesman | Belgium | W.-Vlaanderen |
| 146030 | Julien Rochefort | France | Brittany |
| 570183 | Mark Shorten | Ireland | Cork |
| 589122 | Stanislas Wroza | France | Bretagne |
| 594236 | Stanislas Wroza | France | Pas-de-Calais |
| 435099 | Michèle Peron | Germany | Oberbayern |
| 281355 | Peter Boesman | Belgium | Zeebrugge |
| 711247 | Alan Dalton | Sweden | Stockholms lan |
| 589121 | Stanislas Wroza | France | Finistère |
| 393006 | Louis A. Hansen | Denmark | Kroghage |
| 611904 | Alan Dalton | Ireland | Dublin |
| 110353 | Patrik Aberg | Sweden | Vastergotland |

***Alarm* call *A. cervinus***

| XC Recording | Recordist | Country | Location |
|---|---|---|---|
| 85836 | Christoph Bock | Norway | Slettnes |
| 85840 | Christoph Bock | Norway | Varanger |
| 109145 | Stein O. Nilsen | Norway | Hornoya |
| 396698 | Sunny | Russian Federation | Sakha Republic |
| 138996 | Fernand Deroussen | Norway | Finnmark |
| 240712 | Terje Kolaas | Norway | Finnmark |
| 399824 | Stein O. Nilsen | Norway | Vardo |
| 643913 | Martinez Nicolas | France | Alpes-Cote d'Azur |
| 396674 | Sunny | Russian Federation | Sakha Republic |
| 85844 | Christoph Bock | Norway | Norrkin |

***Alarm* call *A. petrosus***

| XC Recording | Recordist | Country | Location |
|---|---|---|---|
| 188716 | Anthony McGeehan | Ireland | County Mayo |
| 435106 | Patrick Franke | Norway | Runde |
| 146033 | Julien Rochefort | France | Brittany |
| 345773 | Tero Linjama | Finland | Uusikaupunki |
| 53806 | Niels Krabbe | Sweden | Vadero |
| 483209 | Grzegorz Lorek | Norway | More og Romsdal |
| 734118 | David Tattersley | United Kingdom | Wales |
| 731687 | Sonotheque ADVL | France | Normandie |
| 666525 | Peter Mattsson | Sweden | Blekinge lan |
| 588514 | Stanislas Wroza | France | Bretagne |

***Alarm* call *A. pratensis***

| XC Recording | Recordist | Country | Location |
|---|---|---|---|
| 107492 | Patrik Aberg | Sweden | Vastergotland |
| 395929 | Livon | Estonia | Saarema |

| | | | |
|---|---|---|---|
| 423325 | Lars Edenius | Sweden | Vasterbottens lan |
| 651319 | Uku Paal | Estonia | Rannu |
| 578647 | Thomas Bergman | Sweden | Harjedalen |
| 595966 | Simon Elliot | United Kingdom | Northumberland |
| 734963 | Irish Wildlife Sounds | Ireland | County Offaly |
| 724391 | Markus Jacobs | Germany | Lower Saxony |
| 740997 | Irish Wildlife Sounds | Ireland | County Wexford |
| 656491 | Uku Paal | Estonia | Roude |
