## Supplemental results for "VOCAL SIMILARITY AND DIFFERENCES IN FOUR SPECIES OF THE GENUS *ANTHUS*: ACOUSTIC FEATURE ANALYSIS OF SOME COMMON CALLS"

### MP3 COMPRESSION EFFECT ON SPECTROGRAM FREQUENCY CONTOUR

Two call types (*tsip* and *soft*) were selected for the test, because they were more frequency modulated and complex than *alarm* call. Therefore they can be more useful to test if Frequency Contour 75 (FC75) function of Raven Pro is affected by MP3 compression at two quality levels. 30 calls (15 *tsip* and 15 *soft*) were randomly selected from 30 different recordings in wav original format. Each call was compressed to one MP3 files of medium quality (170/210 kbps) and one of low quality (145/185 kbps), using Audacity software with variable bitrate encoder. The resulting contours of wav files in original format, MP3 medium quality and MP3 low quality were compared.

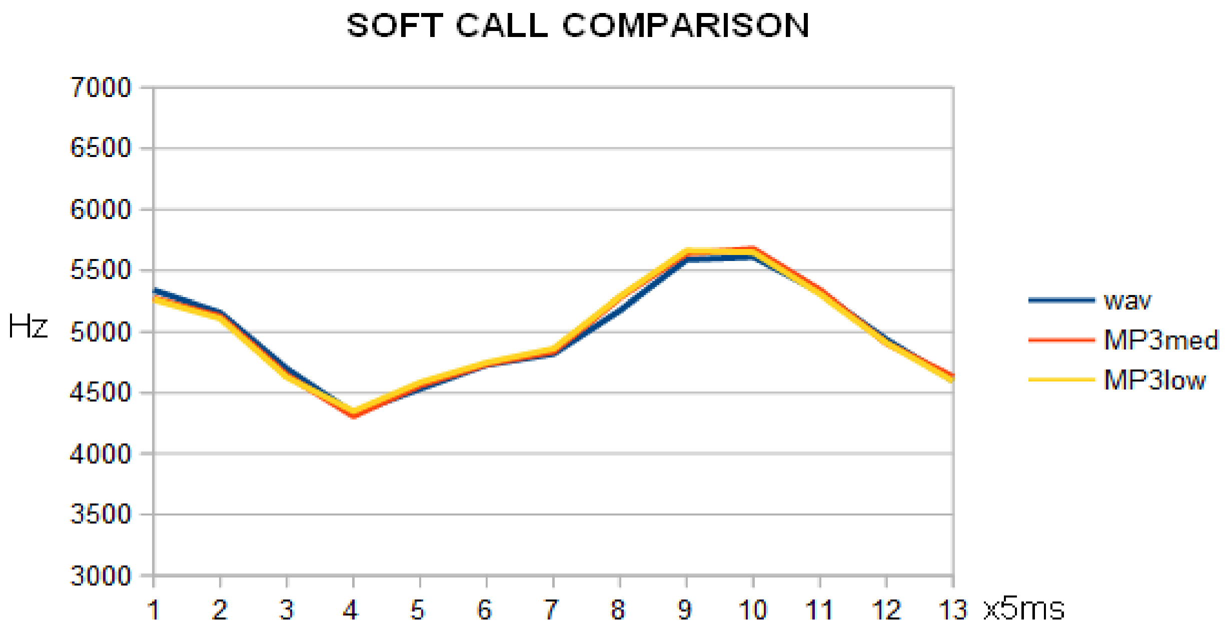


**Figure 1**: Average contour (FC75) of *A. Pratensis soft* call in original uncompressed format (wav), MP3 medium quality (MP3med), MP3 low quality (MP3low). N=15

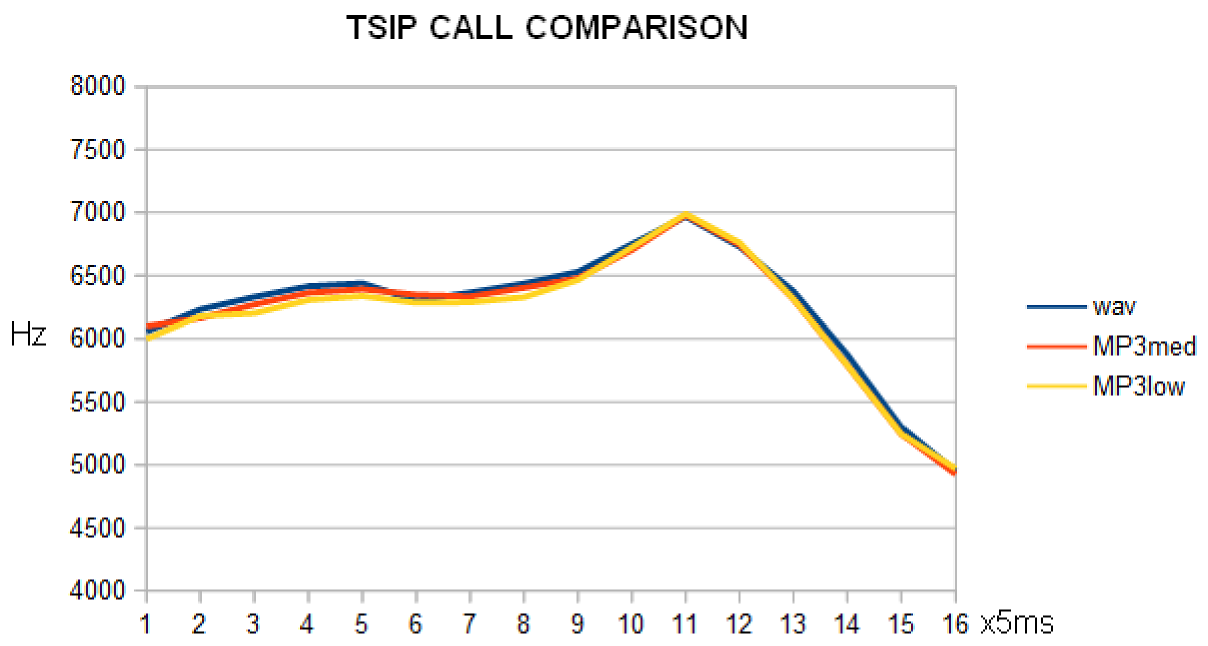


**Figure 2**: Average contour of *A. pratensis tsip* call as in Figure 1. N=15

Cross correlation was calculated (using fcc function of R) between each wav file and its two correspondent MP3 files. Average cross correlation was then calculated for all wav versus all MP3 medium quality files (N=30) and for all wav versus all MP3 low quality files (N=30), yelding the following results: 0,984 +/- 0,023 (wav vs MP3 medium) and 0,979 +/- 0,031 (wav vs MP3 low). The difference between cross correlation for the two quality levels is statistically not significant (T for two sample paired test = 0,50).

In conclusion these results show that MP3 compression does not affect significantly the FC75 parameter and that the general acoustic features of the calls can be represented in a sufficient accurate way either in original wav or in MP3 compressed format.
